# Single-cell molecular connectomics of intracortically-projecting neurons

**DOI:** 10.1101/378760

**Authors:** Esther Klingler, Julien Prados, Justus M Kebschull, Alexandre Dayer, Anthony M Zador, Denis Jabaudon

## Abstract

The neocortex is organized into distinct areas, whose interconnectivity underlies sensorimotor transformations and integration^1–7^. These behaviorally critical functions are mediated by intracortically-projecting neurons (ICPN), which are a heterogeneous population of cells sending axonal branches to distinct cortical areas as well as to subcortical targets^8–10^. Although population-based^11–14^ and single-cell^15–19^ intracortical wiring diagrams are being identified, the transcriptional signatures corresponding to single-cell axonal projections of ICPN to multiple sites remain unknown. To address this question, we developed a high-throughput approach, “Connect_ID_”, to link connectome and transcriptome in single neurons. Connect_ID_ combines MAPseq projection mapping^17,20^ (to identify single-neuron multiplex projections) with single-cell RNA sequencing (to identify corresponding gene expression). Using primary somatosensory cortex (S1) ICPN as proof-of-principle neurons, we identify three cardinal targets: (1) the primary motor cortex (M1), (2) the secondary somatosensory cortex (S2) and (3) subcortical targets (Sub). Using Connect_ID_, we identify transcriptional modules whose combined activities reflect multiplex projections to these cardinal targets. Based on these findings, we propose that the combinatorial activity of connectivity-defined transcriptional modules serves as a generic molecular mechanism to create diverse axonal projection patterns within and across neuronal cell types.

---

The primary somatosensory cortex (S1) is strongly connected with M1 and S2 *via* distinct functional pathways^2,3^. While M1- and S2-projecting ICPN have largely mutually exclusive projections to these two areas, how this specificity emerges postnatally and whether projections to other targets (including subcortical targets) exist has not been systematically examined. To address this question, we used MAPseq to identify the projections of single neurons in S1 (**Fig. 1a, Fig. S1a, Supplementary Note 1**)^17,20^. MAPseq allows single-cell reconstruction of axonal projections to multiple remote targets using anterograde transport of a barcoded RNA from the soma^20^.

**Fig. 1:**
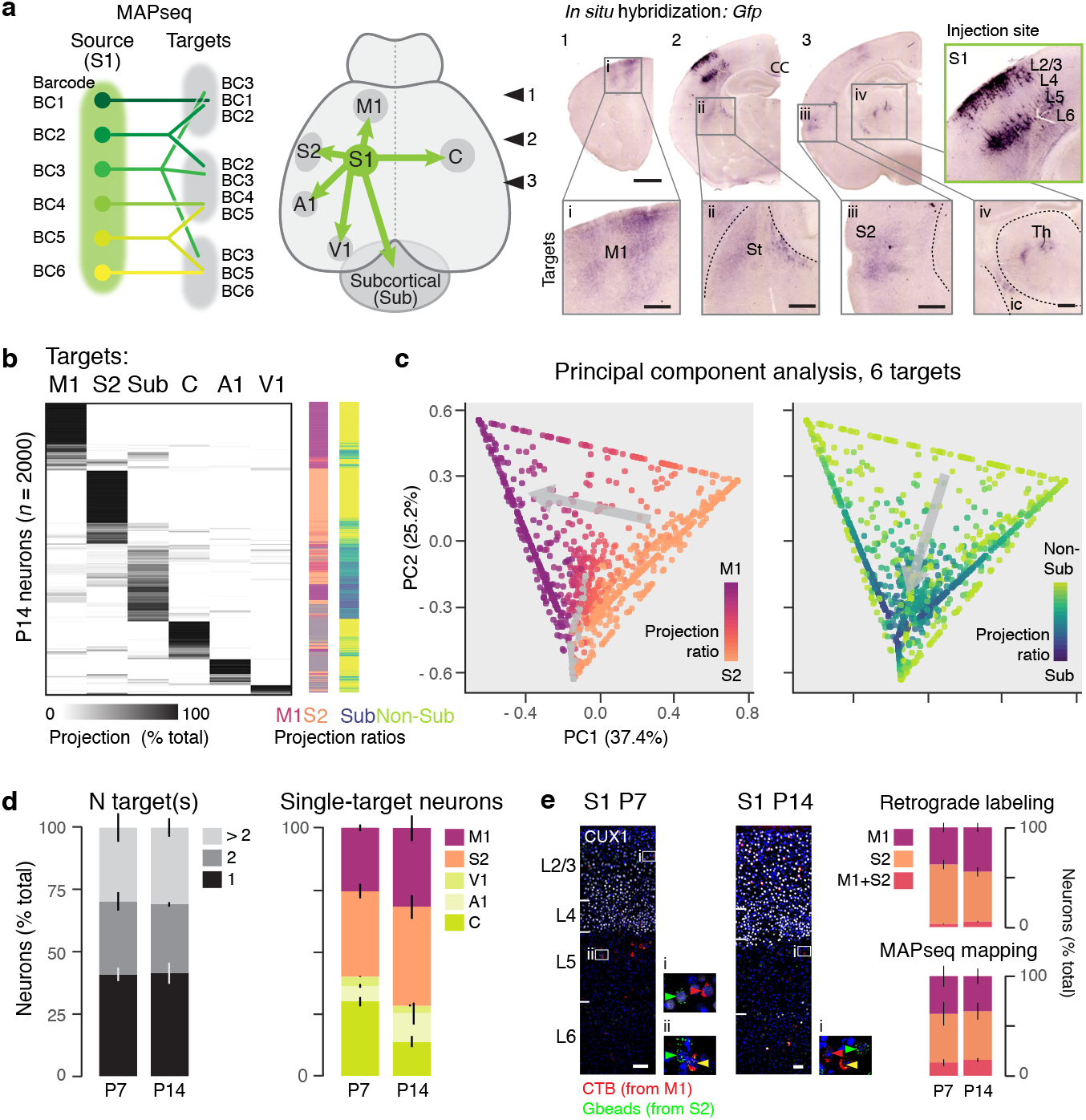
S1 ICPN have two cardinal axonal projection motifs. **a**, Schematic diagram of the experimental approach. MAPseq^20^ is used to map the intracortical and subcortical axonal projections of S1 intracortically-projecting neurons (ICPN). *In situ* hybridization for Gfp showing that barcode-Gfp mRNA is transported from neuronal somas at the injection site (framed in green) into axons at target sites (framed in gray), **b**, Multiplex axonal projection of 2000 single S1 ICPN at P14 (*n*= 5 pups). Total barcode amounts were normalized to 100% for each cell. M1/S2 and Sub/Non-Sub projection ratios are indicated for each neuron, **c**, Principal Component Analysis (PCA) of projections shows that two principal motifs drive ICPN projection diversity: (1) projection to either M1 or S2 (PC1), and (2) projection to either Sub or Non-Sub (*i.e*. all intracortical targets) (PC2). The sharply delineated shape of the PCA reflects the constrained distribution of points to a sum of 1, see also Supplementary Fig. 2b. **d**, The number and distribution of ICPN target(s) is stable between P7 and P14 (number: *P* > 0.99; distribution: *P* > 0.99, two-way ANOVA). **e**, Projections to M1, S2, and both M1 and S2 are stable between P7 and P14, as assessed with retrograde labeling from M1 and S2 (*n* = 3 pups per age) and with MAPseq mapping (*P* = 0.87, two-way ANOVA). Error bars, SE (d, e). Scale bars, 100 μm. S1, primary somatosensory cortex; M1, primary motor cortex; S2, secondary somatosensory cortex; A1, primary auditory cortex; V1, primary visual cortex; St, striatum; Th, thalamus; CC, corpus callosum.

**S**omatosensory input first reaches S1 from where it is forwarded to other cortical and subcortical targets^1–7^. To investigate efferent S1 connectivity with single-cell resolution, we examined singleneuron projections to the following six potentially functionally relevant targets: M1, S2, primary auditory (A1) and visual (V1) cortex, contralateral cortex (C) and Sub (striatum and thalamus) (**Fig. 1a, b**). We performed injections in S1 on either postnatal day (P) 7 (as axons are reaching their targets) or P14 (when projections are largely established)^8,21,22^ and collected tissue from both the injection and target sites (**Fig. S1b, c**, and see Methods).

MAPseq analysis at P14 revealed single ICPN with a variety of projections to target sites (**Fig. 1b**, left). M1, S2, and Sub were the main targets of S1 axons; 87% of ICPN contacted at least one of these sites.

Principal component (PC) analysis of projection targets highlighted two main axes of organization, which we termed projection motifs: projection to either M1 or S2 (PC1) and projection to either Sub or Non-Sub (PC2) (**Fig. 1c** and **Fig. S2b**; Non-Sub referring to all intracortical targets). A spectrum of projection strengths was present within each of these two motifs, with some neurons projecting strongly to one target and others having more balanced projections to multiple targets (as calculated using M1 / (M1 + S2) and Sub / (Sub + Non-Sub) projection ratios; **Fig. 1b**, right, and **c**). These two projection motifs were independent of each other: projection ratio within one motif was not predictive of the projection ratio within the other (**Fig. S2c**; Pearson correlation coefficient: *r* = 0.02), consistent with the results of the PC analysis above. These findings thus identify two independent projection motifs underlying S1 ICPN wiring diversity.

Most neurons had only one or two axonal targets (**Fig. 1d**); when examined at P7, connectivity was similar to that found at P14, suggesting directed projections to target sites as opposed to exuberant growth followed by axonal retraction (*P* > 0.99, two-way ANOVA; **Fig. 1d, Fig. S2**). Accordingly, the small fraction of neurons projecting to both M1 and S2 was unchanged between P7 and P14 (*P* > 0.99, two-way ANOVA), which we confirmed using dual retrograde labeling with classical dyes (**Fig. 1e**). This suggests that M1- and S2-projecting neurons do not emerge from an initial population of dual-projecting ICPN, but instead are specified early on. Together, these data indicate that S1 ICPN have a mostly sparse and directed connectivity during the second postnatal week, and exclude widespread pruning as a mechanism to sculpt intracortical connections.

Given their prevalence and functional importance, we focused on the projection diversity of M1-projecting neurons and S2-projecting neurons, which represent ~70% of S1 ICPN at P14. We used hierarchical clustering to classify projection patterns (**Fig. 2**, see Methods). This analysis revealed that projection to M1 or S2 represents a primordial segregation of S1 ICPN, but that beyond this dichotomy, M1- and S2-projecting neurons have essentially identical additional targets, as revealed by a matching distribution into seven main projection patterns (**Fig. 2**). More than half of the neurons projected exclusively to M1 or to S2 (63% in the case of M1-projecting neurons; 59% for S2-projecting neurons; Pattern 1) and a quarter of all neurons had subcortical projections (28% of M1-projecting neurons; 27% of S2-projecting neurons; Patterns 2 and 3). Additional projection patterns included dual projections to either M1 or to S2, along with projections to the contralateral cortex or to A1 or to V1 (to C: 7%, Pattern 4; to A1: 3.5%, Pattern 5; to V1: 2.5%, Pattern 6). Altogether, M1-only, S2-only, M1 + Sub and S2 + Sub projecting neurons accounted for 87% of all S1 M1- and S2-projecting neurons, in line with data obtained using single-cell anterograde labeling^18^. Highlighting the importance of these targets in defining projection diversity, a projection-based PC analysis revealed M1 *vs*. S2 and Sub *vs*. Non-Sub as the two driving motifs of this hierarchical clustering (**Fig. 2b**).

**Fig. 2:**
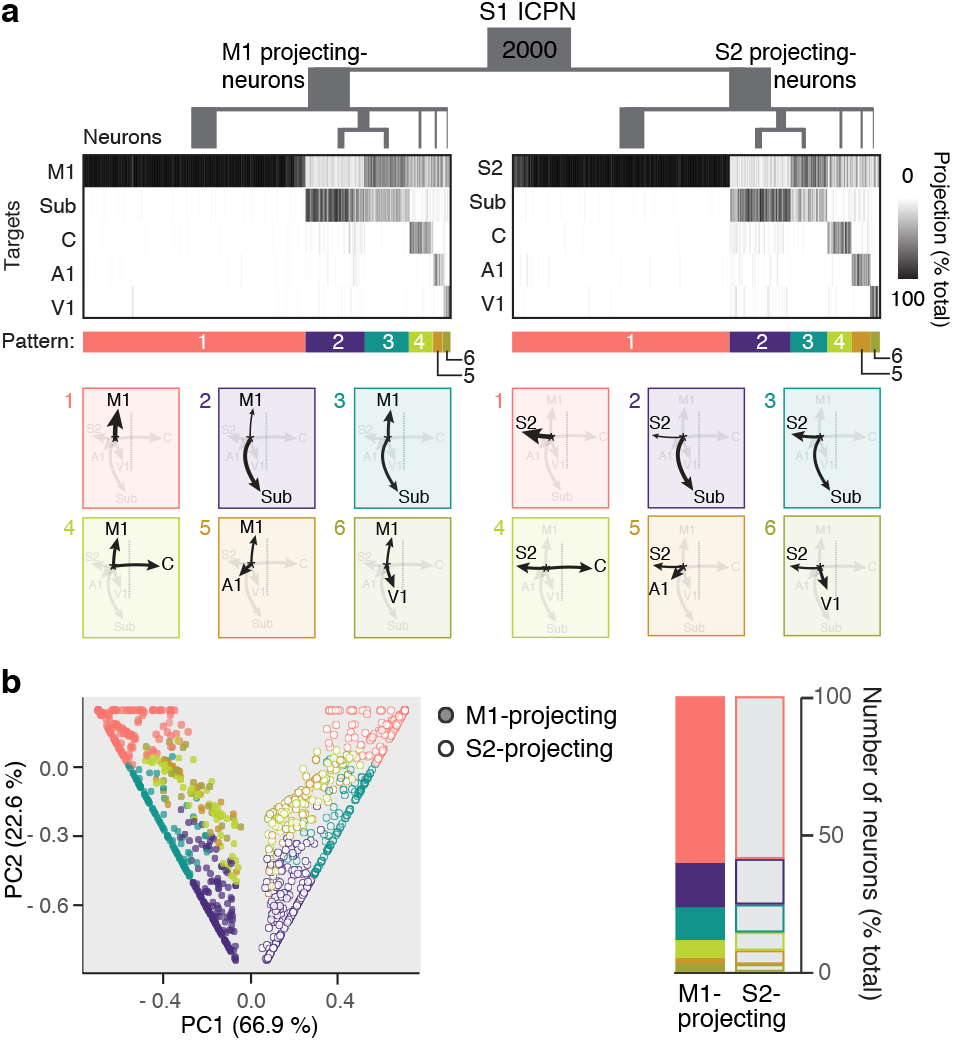
M1- and S2-projecting ICPN have otherwise identical projection patterns. **a**, Cluster analysis of M1-or S2-projecting ICPN at P14 revealing 6 projection patterns (*n* = 1000 M1 or S2 ICPN, Manhattan distances, see Methods). Note the similar distribution into patterns 1-6 for M1- and S2-projecting ICPN. **b**, PCA (left) and distribution of P14 M1-or S2-projecting ICPN projection patterns (right; *P* = 0.73, χ2 test). The symmetrical layout of M1- and S2-projecting ICPN in the PCA reflects similar projection patterns. The sharply delineated shape of the PCA reflects the constrained distribution of points to a sum of 1.

We next investigated the transcriptional signatures corresponding to the multiplex projections of S1 ICPN. For this purpose, we combined MAPseq mapping^17,20^ with single-cell RNA sequencing of neurons at the injection site to link the transcriptional identity of barcode-identified neurons with their corresponding projections to our targets of interest (“Connect_ID_”, *n* = 161 quality-controlled neurons at P14; **Fig. 3a, Fig. S3, Table S1** and **Supplementary Note 2**). In a subset of experiments, we microdissected superficial and deep layers and generated a training set to define laminar identity (**Fig. S4** and Methods). MAPseq analysis of the Connect_ID_ dataset confirmed the distribution of axonal projection patterns reported above (**Fig 3b**, left). However, a corresponding transcriptional organization was not readily apparent using classical analytical strategies such as k-means clustering and tSNE dimensionality reduction (**Fig. 3b**, right, cells aligned as in **3b**, left, and data not shown). Accordingly, while PC analysis of MAPseq projections revealed the two projection motifs described earlier (*i.e*. M1 *vs*. S2 and Sub *vs*. Non-Sub) (**Fig. 3c**, top), transcriptome-based analysis only showed a layer-related distribution (**Fig. 3c**, bottom).

**Fig. 3:**
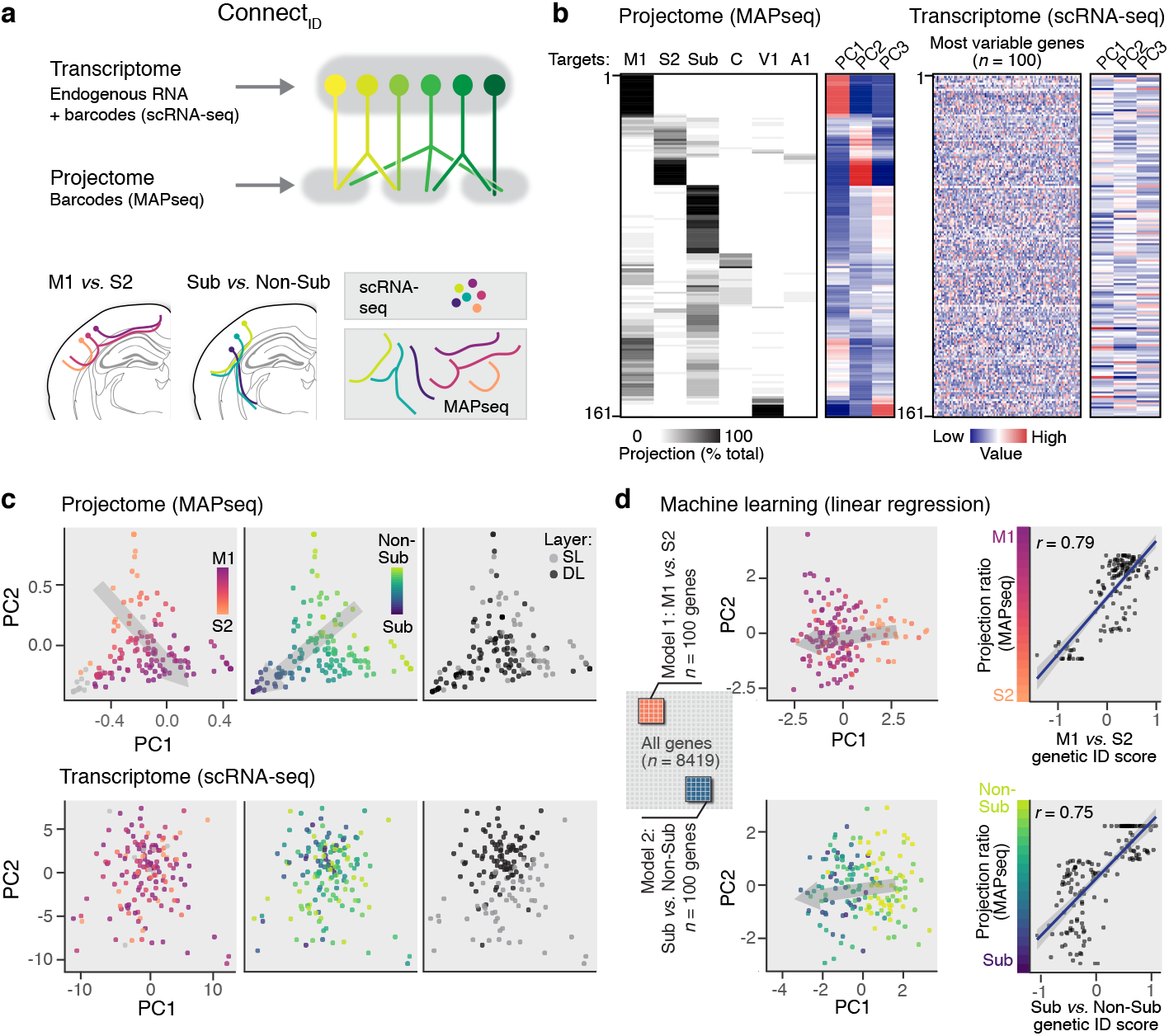
Connect_ID_ links connectome with transcriptome in single neurons. **a**, Schematic diagram of the Connect_ID_ strategy (top) and of the ICPN subtypes studied here (bottom). **b**, Multiplex axonal projection of 161 S1 neurons (left) with corresponding gene expression (right). Projectome-based structure is not readily visible in the transcriptomic data. **c**, While principal component (PC) analysis of anatomical projection patterns identifies the M1 *vs*. S2 and Sub *vs*. Non-Sub projection motifs, only a laminar location-related organization is visible in the transcriptomic data. **d**, Machine-learning approach using M1 *vs*. S2 and Sub *vs*. Non-Sub projecting neurons as training sets identifies a core set of genes predicting anatomical projection ratios within these two motifs.

Since global transcriptional activity did not evidently reflect neuronal projections, we developed a machine learning approach to isolate subsets of genes associated with these two projection motifs^23^. We generated two linear regression models with M1-*vs*. S2-projecting neuron transcripts ([M1 *vs*. S2] model) and Sub-*vs*. Non-Sub-projecting neuron transcripts ([Sub *vs*. Non-Sub] model) to identify the genes with the highest weight in distinguishing model-respective projections (*n* = 100 transcripts per model; **Fig. S5a** and Methods). The weighted average of gene expression was used to define a transcription-based projection identity score for each neuron (“genetic ID” score). By focusing our analysis on the expression of this core set of genes, we uncovered projection motif-related transcriptional identities (**Fig. 3d, Fig. S5b, Table S2**).

Both models accurately predicted the spectrum of anatomically-defined (*i.e*. MAPseq-mapped) projections within each motif: for example, neurons with a high genetic ID score for M1 *vs*. S2 projections had a high MAPseq-defined M1/S2 projection ratio. Similarly, Sub *vs*. Non-Sub projections were correctly identified (**Fig. 3d and Fig. S5a**). Predictions of M1 *vs*. S2-projecting identity were less accurate than for Sub-*vs*. Non-Sub-projections, suggesting that subcortical projections are associated with more salient transcriptional features than distinct intracortical connections (**Fig. S5a**)^24^. Predictions of the [M1 *vs*. S2] model were independent of the laminar position of neurons, consistent with the isotropic distribution of M1- and S2-projecting neurons in superficial and deep layers. In contrast, in the [Sub *vs*. Non-Sub] model, subcortical projections were correlated with deep layer position, in line with the preferential location of subcortically-projecting neurons^25^ (**Fig. S4 and S5**).

Consistent with the independence between M1/S2 and Sub/Non-Sub projection motifs (see above), the genesets in the [M1 *vs*. S2] model and the [Sub *vs*. Non-Sub] model did not overlap (**Table S2**). Enriched ontologies included neuronal projection (e.g. *Ndrg2, Llcam*), and synaptic functions (*e.g. Chrm1, Homer2, Pkp4, and Cdh8*) (**Fig. S5b, c**). In the [Sub *vs*. Non-Sub] model lamina-enriched transcripts included *Cux2* and *Pou3f2*. Laminarly-enriched transcripts only formed a subset of the geneset of this model, since only 12/100 genes were shared with the data generated from the superficial *vs*. deep layer training set and only 4/9 genes referenced in the Allen Brain Atlas developmental database had lamina-specific expression (**Fig. S5a, d, Table S2**). Altogether, these data identify three core projection motif-based transcriptional signatures of S1 ICPN corresponding to M1 projections, S2 projections, and Sub projections.

To investigate multiplex projections, we combined the two aforementioned models. Individual neurons were displayed based on their genetic ID scores in the [M1 *vs*. S2] model and in the [Sub *vs*. Non-Sub] model (**Fig. 4a**). This genetically defines their combined projections to M1, S2 and Sub, and creates a two-dimensional axonal projection space within which single and combined gene expression patterns can be examined. In some cases, single transcripts were associated with specific projection patterns. For example, the transcription factor *Lhx2*, which is required for barrel cortex formation^26^ was enriched in M1-projecting neurons, and *Homer2*, which regulates metabotropic glutamate receptor function, was enriched in S2-projecting neurons, consistent with the important role of metabotropic transmission in this cortical area^27^. In general however, individual genes only weakly defined multiplex projections (**Fig. 4b**). In contrast, we found a close correspondence with the overall expression of genes within each of the 3 core genesets identified above (**Fig. 4c**). Thus, projection patterns are associated with the cumulative expression of genes within connectivity-defined transcriptional modules rather than with the expression of single master genes.

**Fig. 4:**
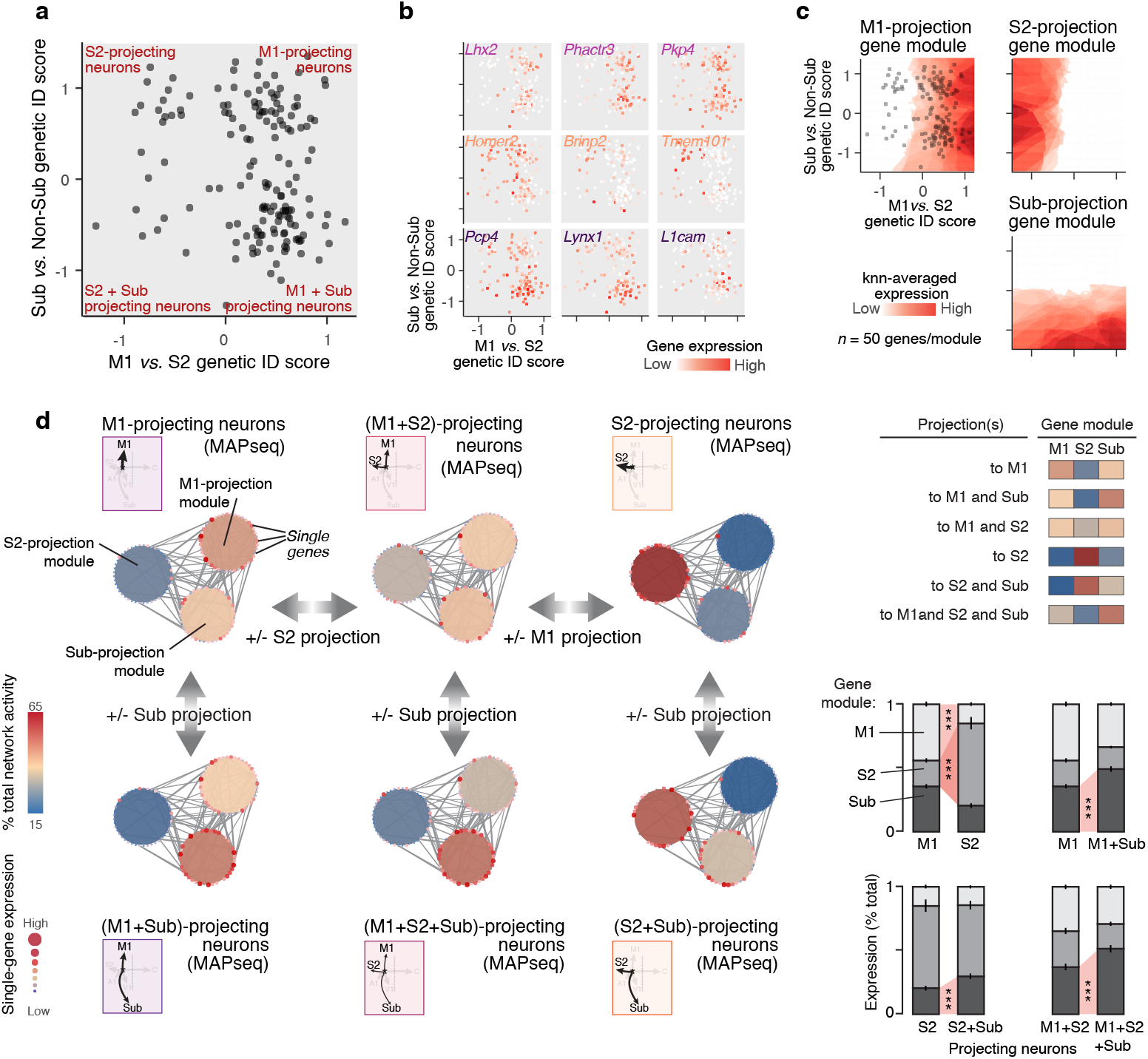
Activity of target-specific transcriptional modules defines the multiplex projections of ICPN. **a**, 2D-map showing distribution of single neurons with respect to their genetically-defined projections to M1, S2 and Sub targets, obtained by combining the two models presented in Fig. 3. **b**, Expression of select genes using the map generated in a. Distinct target-related gene expression patterns are visible. **c**, Expression of the core M1, S2 and Sub target genes identified in Fig. 3 outlines three projection-related gene expression modules (projection module). **d**, Combinatorial activity of the M1, S2 and Sub modules within neurons with anatomically defined (*i.e*. MAPseq-mapped) projections. For example, in ICPN with both M1 and Sub projections, the Sub projection module is more active than in neurons projecting only to M1. Lines between single genes indicate known functional protein-protein relationships (STRING database^40^). Top right panel: summary; same color code as left. Bottom right panel: quantification; two-way ANOVA; Error bars, SE.

Finally, we sought for the transcriptional logic underlying simultaneous projections to M1, S2 and Sub. We hypothesized that projections to multiple targets might involve several individual connectivity-defined transcriptional modules in single neurons. For example, the transcriptional signature of ICPN projecting both to M1 and Sub might consist in a combination of M1 and Sub genetic identities. We examined the combinatorial transcriptional activity of these three modules in neurons with anatomically defined (*i.e*. MAPseq-mapped) projections. Functional protein association network analysis^28^ revealed multiple interactions which could account for cross-talk within and across modules. Focusing on six archetypical projection combinations in MAPseq-mapped neurons (*i.e*. M1-only, S2-only, M1+Sub, S2+Sub, M1+S2 and M1+S2+Sub), we reveal that combinatorial activity of individual gene modules indeed reflects combinatorial projections to corresponding targets (**Fig. 4d**).

Together, our data establish a first high-throughput correspondence between connection-based identity and transcriptional identity in single neurons, allowing identification of gene modules whose combined activity defines axonal projection motifs. These modules may represent downstream targets of master, terminal-selector-type genes, which are transiently expressed earlier in development and drive target-specific developmental programs^29–34^. Alternatively, the transcriptional networks outlined here may not be directly related to developmental events, but instead reflect transcriptional states associated with the constraints of specific connectivities. Based on protein network analysis, the identified gene modules are reciprocally functionally interlinked, such that cross-regulation of network activity may involve processes such as cis/trans activity of additional genes as well as complementary/mutually exclusive 3D chromatin conformations and associated epigenetic events^35^.

The genetic organization identified here provides a parsimonious mechanism to generate a spectrum of projections within genetically-defined classes of neurons as well as a means to create new projection patterns through cross-regulation of multiple, discrete, gene modules. Such combinatorial coding of projection patterns may have been recruited during evolution to increase neuronal diversity and generate new wiring diagrams from an initially limited set of cellular elements.

## Acknowledgements

We thank the Genomics Platform and FACS Facility of the University of Geneva, A. Benoit for technical assistance; and members of the Jabaudon laboratory, as well as G. Limoni, U. Tomasello, and E. Azim for constructive comments on the manuscript. The Jabaudon Laboratory is supported by the Swiss National Science Foundation and the Brain and Behavior foundation: E.K. is supported by a grant from the Machaon Foundation. J.P. is supported by the Public Instruction Department, Geneva. A.D. is supported by the Swiss National Foundation Synapsy (grant 51NF40-158776). The Zador Laboratory is supported by the following funding sources: National Institutes of Health [5RO1NS073129 to A.M.Z., 5RO1DA036913 to A.M.Z.]; Brain Research Foundation [BRF-SIA-2014-03 to A.M.Z.]; IARPA MICrONS [D16PC0008 to A.M.Z.]; Simons Foundation [382793/SIMONS to A.M.Z.]; Paul Allen Distinguished Investigator Award [to A.M.Z.].

## Author contributions

E.K., and D.J. conceived the project and designed the experiments. E.K. performed the experiments. E.K. and J.P performed the bioinformatics analyses. J.M.K. and A.M.Z provided Sindbis virus and shared expertise on MAPseq. E.K. and D.J. wrote the manuscript. J.M.K., A.M.Z, and A.D. revised and edited the manuscript.

## Materials and methods

### Mouse strains

C57Bl6/J male and female pups from Charles River Laboratory were used. Experiments were carried out in accordance with permission of the Geneva cantonal authorities.

### Stereotaxic injections

Anesthetized pups were placed in a stereotaxic apparatus on either postnatal day (P) 7 or 14 and were injected with barcoded Sindbis virus (80 nl)^20^, or red Retrobeads™ IX and green Retrobeads™ from Lumafluor (100 nl) (**Fig. 1e**), or Alexa-488 conjugated cholera toxin subunit B (CTB, Invitrogen, #C-34775) (100 nl) (**Fig. 1e**).

Coordinates of injection sites from lambda, along antero-posterior (AP) and along medio-lateral (ML) axes:

- M1 injection site: P7, AP: 3.3, ML: 1.3; P14, AP: 4.5, ML: 1.5.
- S2 injection site: P7, AP: 3, ML: 4.2; P14, AP: 3.2, ML: 4.5.
- S1 injection site: P7, AP: 3, ML: 3; P14, AP: 3.2, ML: 3.2.

For the initial mapping experiments (Figs. 1 and 2) Sindbis virus-injected pups were collected at either 14 hours (*n* = 2 pups at P7; *n* = 3 pups at P14) or 24 hours (*n* = 2 pups per age) post-infection, when Sindbis *Gfp* RNA is present in axons (**Fig. 1b; Supplementary Note 1**). This time point allows for preservation of the integrity of the infected cells (and thus access to transcriptomic identities) without affecting the efficiency of MAPseq mapping (see **Supplementary Note 1**). For Connect_ID_ experiments, pups were collected 14 hours post-infection. Retrobeads/CTB injected pups were collected 48 hours post-injection.

### Immunohistochemistry

Postnatal mice were perfused with 4% paraformaldehyde (PFA) and brains were fixed overnight in 4% PFA at 4 °C. Eighty postnatal mice were perfused with 4% paraformaldehyde (PFA) and pre-incubated 2 h at room temperature in a blocking/permeabilizing solution containing 5% bovine serum albumin and 0.3% triton X-100 in PBS, and incubated for 2 days with primary antibodies at 4°C. Sections were then rinsed 3 times in PBS and incubated with the corresponding Alexa-conjugated secondary antibodies (1:500; Invitrogen) for 2 h at room temperature. Primary antibodies and their dilutions were: chicken anti-GFP (Invitrogen, #A10262, 1:200), rabbit anti-CUX1 (Santa Cruz, #SC-133606, 1:250).

### *In situ* hybridization

For antisense *Gfp* probe synthesis, DIG-labeled antisense RNA probe was obtained after *in vitro* transcription of GFP brain expressing-transgenic mouse line cDNA using specific primers (forward primer: CCATCCTGGTCGAGCTGG; T7Reverse primer: CGATGTTAATACGACTCACTATAGGGCTTCTCGTTGGGGTCTTTGC). *In situ* hybridization on slides was performed according to methods described previously^36^. In brief, hybridization was carried out overnight at 60°C with the digoxigenin (DIG)-labelled Gfp RNA probe. After hybridization, sections were washed and incubated with alkaline phosphatase-conjugated anti-DIG antibody (Roche, #11093274910, 1:2000) for 2 days at 4°C. After incubation, sections were washed and the color reaction was carried out overnight at room temperature in a solution containing NBT (nitro-blue tetrazolium chloride) and BCIP (5-brom-4-chloro-3’-indoly phosphate *p*-toluidine salt) (Roche, #000000011681451001). After color revelation, sections were washed, post-fixed for 30 min in 4% PFA and mounted with Fluoromount (Sigma).

### Image acquisition and quantifications

All images from immunohistochemistry were acquired on a Nikon A1r spectral confocal microscope, equipped with 20x 0.5 CFI Plan Fluor WD objective. All images from *in situ* hybridizations were acquired on a Zeiss Axioscan.Z1 slide scanner, equipped with a 10x/NA 0.45 Plan Apochromat objective, and a Hitachi HV-F202FCL camera.

### Tissue microdissection, cell sorting and single-cell RNA sequencing

*MAPseq:* Acute coronal brain sections were cut on a vibrating microtome (Leica, VT1000S) and brain regions were microdissected with micro-scalpel using a Leica Dissecting Microscope (Leica, M165FC) in ice-cold oxygenated artificial cerebrospinal fluid (ACSF) under RNase-free conditions. Brain from distinct pup was microdissected separately, on ice.

P7: *n* = 4 pups; *n* = 1 litters; sections thickness = 600 μm.

P14: *n* = 5 pups; *n* = 2 litters; sections thickness = 600 μm.

Microdissected brain tissues were collected in TRIzol reagent-containing tubes (ThermoFisher, #10296-010), mechanically dissociated and immediately stored at −80°C. Throughout the procedure, sample cross-contamination was carefully avoided. Dissected samples were processed for sequencing as previously described^17,20^. Samples were mixed with spike-in RNA and performed reverse transcription, production of double-stranded cDNA, treatment with ExonucleaseI (NEB), and two rounds of nested PCR using primers 5′-CTCGGCATGGACGAGCTGTA-3′ and 5′-CAAGCAGAAGACGGCATACGAGATCGTGATGTGACTGGAGTTCCTTGGCACCCGAGAATTCCA-3 7 for the first PCR and primers 5′-AATGATACGGCGACCACCGA-3′ and 5′-CAAGCAGAAGACGGCATACGA-3′ for the second PCR using Accuprime Pfx polymerase (ThermoFisher, #12344-040) as previously described^17,20^. Finally, the resulting PCR amplicons were gel-extracted using Qiagen MinElute Gel extraction kit according to the manufacturer’s instructions and the cDNA purified with magnetic Agencourt AMPureXP beads (Beckman, #A63881). Finally, we sequenced the library on an Illumina HiSeq4000 next gen sequencer using SBS kit (Illumina) pool of primers (HP10, HP12) for single-end 106 base-pair sequencing.

*Single-cell RNA sequencing*: the primary somatosensory cortex was microdissected as aforementioned. Cells were further dissociated by incubating micro-dissected samples in 0.5 mg/mL pronase (Sigma, #P5147) at 37°C for 10 minutes, followed by incubation in 5% bovine serum albumin for 3 minutes and manual trituration in ACSF using pulled glass pipettes. Cells were then centrifuged for 10 minutes at 600 rpm and resuspended before filtration using a 70 mm cell strainer (ClearLine, #141379C). Cells were then incubated for 10 minutes at 37°C with Hoechst (0.1 mg/mL) and isolated using a Beckman Coulter Moflo Astrios FAC-sorter. Singlet Hoechst^+^ cells were sorted according to their Forward and Slide scattering properties, and their negativity for Draq7TM (Viability dye, Dar red DNA intercalating agent, Beckman Coulter, #B25595). 5000 to 10’000 cells were FAC-sorted for each experiment. Three microliters of C1 Suspension Reagent (Fluidigm) was added to 10 μl of FACsorted cells, which were captured into AutoPrep integrated fluidic circuit (IFC) designed for 10 to 17 μm diameter-cells (Fluidigm, #100-57-80).

RNA was extracted using an RNeasy kit (Qiagen, #74034), and quality control was done using 2001 Bioanalyzer from Agilent. cDNA libraries were obtained using SMARTseq v4 kit (Clontech, # 634888) and sequenced using HiSeq 2500 sequencer.

### Analyses

All bioinformatics analyses were performed using R programming language and Bioconductor packages. *MAPseq mapping of projection patterns* Reads of the FASTQ files were de-multiplexed by pup and region according to Sindbis index (read position 89-94bp) with 0 mismatch tolerance against the expected target sequences. Reads with an N in the UMI (read position 95-106bp) or in the barcode (read position 1-32bp) sequence were filtered out, and barcodes with identical UMI sequence were kept only once. Then, barcodes were considered “spike” if their tails (positions 25-32bp) match with the sequence “ATCAGTCA” allowing for 1 mismatch; or considered “viral” if the 2bp tail (positions 31-32bp) match “YY”.

Viral barcodes from S1 injection site were used to build a reference library of barcodes for each pup. To correct for sequencing errors, S1 barcodes were mapped on themselves with bowtie v1.1.1 allowing for 3 mismatches. A graph of sequenced S1 barcodes was generated from the mapping result so that the node represented a barcode sequence, and edges linked two barcodes that differ by less than 3 mismatches. To identify barcodes with sequencing errors, the maximal weakly connected components of the graph were calculated with R package igraph. For each component, the most abundant sequence was kept as the error-corrected barcode sequence and the UMI counts were summed up. Error-corrected barcodes found in S1 injection site were checked against the known catalog of barcode for barcoded Sindbis virus. Barcodes found in the target regions were mapped on this S1 reference library with bowtie v1.1.1, allowing for 3 mismatches. At the end of this procedure, barcodes sequenced in the target regions were associated to barcodes sequenced in S1 and thus establish a picture of S1 multiplex projections. The contralateral thalamus (which does not receive input from the cortical injection site) was used as a negative control target. In this region, we found a mean of 4,3 ± 4,9 barcode counts, which we considered as noise value. We therefore excluded barcodes with less than 10 counts in at least one target, as well as those with less than 100 reads in the injection site (S1).

For **Fig. 1** and **Fig. S2**, we randomly selected same number of barcodes per pup *(i.e*. neurons) in order to normalize for potential variability in labeled cell populations due to variability in the depth of injection across cortical layers (P7: *n* = 4; P14: *n* = 5 pups). Projections of single neurons were normalized to 100% and then ordered into heatmaps by their projection similarities. M1 and S2 projecting neurons displayed more than 10% projection in M1 and S2 respectively.

For **Fig. 2**, we randomly selected same number of M1 and S2 projecting neurons (i.e. neurons with more than 10% projection in either M1 or S2, but not in both) and calculated distances between projection patterns using Manhattan (Ward.D2) method to define clusters of projection patterns displaying more than 30% difference within each other and representing more than 1% of the cells. From this first step, we further pooled similar clusters together and ended with 6 clusters of projection patterns. We then checked the distribution of M1/S2 projecting neurons within these projection patterns.

*Connect_ID_: single-cell RNA sequencing combined with MAPseq (see also Supplementary Note 2)*.

Reads were mapped on mouse genome GRCm38 following the same pipeline as in Telley et al. (20 1 6)^37^. Briefly, read1, which contains the UMI sequence, was appended at the end of read2 header. Read2 were further mapped to the mouse genome with Tophat v2.0.13. Resulting alignment files in BAM format were processed with umi_tools^38^ to deduplicate reads with identical UMI. Gene expression quantification was performed with R using summarizeOverlaps method of package GenomicAlignments. Only reads falling into exonic part of a gene are quantified, and this includes 5’ and 3’ UTRs.

We additionally associated to every transcriptome the result of a manual brightfield picture annotation, where the operator checked for the presence of a single cell in the wells of the fluidigm HT800 chips. Only wells where a single cell was observed were kept for further analysis (wells with no cell, cell(s) with convoluted shapes, multiple cells, or cell(s) with debris were excluded).

Reads that did not map on the mouse genome were aligned against Sindbis virus sequence with BWA^39^ for quantification. In addition, unmapped reads were processed with a custom R script to identify the Sindbis barcode sequence of the cell. Using trimLRpatterns method of Biostrings package, this script looked in unmapped reads for the two sequences surrounding the 32bp random barcode of Sindbis virus. If both sequences matched a read with 5% mismatch tolerance and are separated by exactly 32bp, the sequence in between was considered a barcode of a Sindbis-infected cell. Single-cell barcodes identified in S1-cells of a given pup were further corrected for sequencing error following a similar pipeline as described above for MAPseq. Additionally, to ensure that the same barcode has not been sequenced for several single cells, we required that it was at least 3 times more abundant in its associated cell than in any other cell. At the end of the procedure, we obtained the reference library of barcodes of S1 injection sites. This was used to map barcodes found in the target regions and thus to infer connectivity from single-cell transcriptomes. For single cells with multiple barcodes (*n* = 52/174 cells), we measured the distance between distinct barcode profiles (see **Supplementary Note 1**).

All transcriptomic analyses were performed on reads per million (RPM) normalized gene expressions. Before analyzing the transcriptomes of single-cells, we removed cells with more than 30% of mitochondrial RNA or more than 50% of viral reads compared to total mapped reads. We then filtered genes with ontologies related to response to viral infection (using QuickGO from EMBL-EBI Hinxton database), that were enriched when comparing cells with increasing viral load (*i.e*. viral reads) using ordinal regression (**Fig. S3f, Table S1**, and **Supplementary Note 2**). We further corrected all gene expressions for viral load and number of expressed genes^40^ (see **Supplementary Note 2**). We did all analyses on these *n* = 161 cells and *n* = 8419 gene corrected expressions.

For **Fig. S4** and **Fig. 3**, we used machine learning approach: a logistic regression model with regularization was used to build binary prediction models of: superficial (SL) *vs*. deep (DL) layers, Sub *vs*. Non-Sub projecting neurons, and M1 *vs*. S2 projecting neurons. This implementation was provided by bmrm R package (@Manual{, title = {bmrm: Bundle Methods for Regularized Risk Minimization Package}, author = {Julien Prados}, year = {2018}, note = {R package version 3.10},}). We limited the linear model to the top- and bottom-25 genes with highest absolute weight (feature selection) and re-trained a new model on these selected genes. All model performances were addressed by cross-validations, which gave a prediction value to build receiver operating characteristic (ROC) curves. To check for significance of the model performances, the area under curve (AUC) was compared with the AUC of 10’000 prediction models that randomly assigned labels (*i.e*. SL/DL: *P* < 0.0001, Sub/Non-Sub: *P* < 0.0001, M1/S2: *P* = 0.0001).

For **Fig. S5**, heatmaps were performed using the mean expression of each gene per projection neuron class. Gene ontologies were performed on these genesets using GSEA^41^. *In situ* hybridizations at P14 are from the Allen Brain Developmental Mouse Brain Atlas (http://developingmouse.brain-map.org/).

For **Fig. 4**, we took the cell locations in **Fig. 4a** to infer the expression of single genes (**Fig. 4b**) or of each transcriptional module genes by averaging their expression in the 3-nearest neighbor cells (knn average). String analyses were performed using © STRING CONSORTIUM 2018^28^ with the by default setting for confidence (0.4), and further displayed using Cytoscape. Thickness of the lines shows the level of confidence for each interaction.

## Supplementary notes

### Supplementary Note 1. Sequencing of MAPseq barcodes

MAPseq mapping has been established as a reliable method to measure the projection strength of single neurons to multiple targets. The infection time used in previous studies is 44 hours^17,20^. Here, we use shorter infection times (14-24 hours) to minimize the effect of viral infection on endogenous gene expression (see **Supplementary Note 2**). The following experimental observations support that 14 hours is sufficient for barcodes to be present in (even remote) targets, and reliably reflect anatomical connectivity:

1. **Consistency with data obtained with single-cell filling**. A recent study used single-cell filling to measure projections from S1 L2/3 to other targets^18^. The authors report that 3/15 cells projected both to either M1 or S2 and the striatum (20%; our data: 27%), 3/15 projected both to M1 and S2 (20%; our data: 16%), and 1/15 projected to M1+S2+striatum (6.6%, our data: 11%). The data presented here are thus consistent with these values (*P* = 0.11, *χ*^2^-test).
2. **Validation with retrograde labeling**. The data in Fig. 1e, in which retrograde labeling from S2 and M1 is compared to MAPseq mapping show equivalent distributions of M1-projecting and S2-projecting neurons (M1-projecting neurons: *P* = 0.99, S2-projecting neurons: *P* = 0.78, two-way ANOVA). Of note, double retrograde labeling does not appear to be as efficient as MAPseq mapping to label dual-projection neurons (16% with MAPseq vs. 4% with retrograde labeling). This is consistent with the fact that the axon of single neurons needs to be targeted at each of the two injection sites in the case of double-retrograde labeling. The same effect is observed when comparing single-cell filling (i.e. anterograde labeling) with retrograde labeling from multiple targets: while 20% of S1 neurons (3/15) were found to project both to M1 and S2 using single-cell anterograde labeling^18^, less than 2% of S1 neurons had this projection pattern using retrograde labeling^3^.
3. **Equivalent barcode numbers and counts when comparing 14 and 24 hours infection times**. The number of different barcodes and the counts for each barcode at target sites was similar at 24 compared to 14 hours, showing that viral load does not significantly increase between these time points. Number of different barcodes across IC targets: 14hrs, 1060 ± 311; 24hrs, 1340 ± 399; *P* = 0.25, two-way ANOVA. Mean counts per barcode across IC targets: 14hrs, 6280 ± 1500; 24hrs, 8820 ± 3050; *P* = 0.28, two-way ANOVA. These values were also constant when examining the distinct targets individually, including for the most remote one, *i.e*. the contralateral cortex (P > 0.99 for both number and counts of barcodes, two-way ANOVA).
4. **Independence between barcode counts and distance to target site**. The closest target of S1 ICPN is S2 and the furthest targets are M1 and C (compared to the S1:S2 distance, S1:M1 = ~2x and S1:C= ~6x)^18^. Despite this range of distances, the number of barcodes and the barcode counts in each target is not correlated to the distance to S1 (R^2^ < 0.4 for each condition) but rather reflects projection strengths (see 1 and 2, above).
5. **Robust readout of single-neuron projection patterns**. We compared the closest with brain match *vs*. the closest across brain match of all barcode profiles for all pairs of pups at P14 (see ref. 20 for details). The shift in the within-pups reflects the higher fraction of closely matched projection profiles both after 14 and 24 hours of infection, confirming the reliability of MAPseq in both conditions^20^.

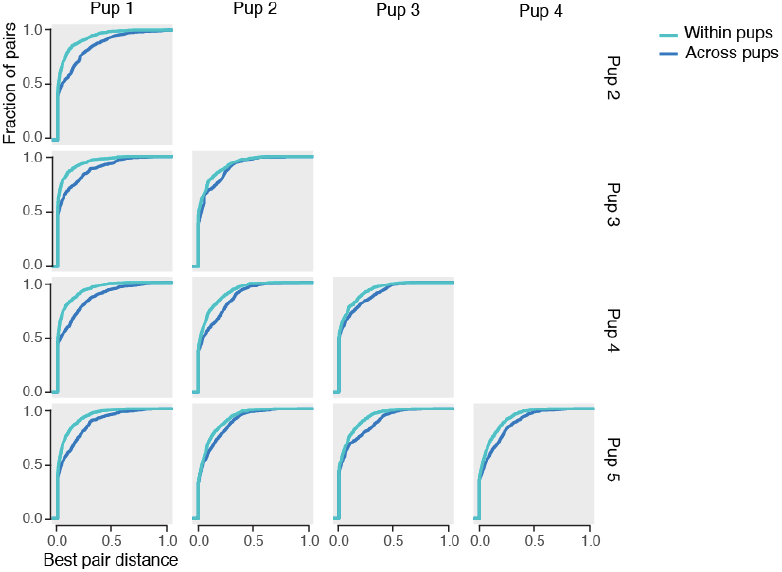
Cumulative distribution of distances between the best barcode pairs within pup and across pups. The shift in the within-animal distribution reflects the higher fraction of closely matched projection profiles, consistent with double infection. Infection time: Pups 1-3: 14 h, Pups 4,5: 24 h.
6. **Replicable barcode distributions in single cells containing more than 1 barcode**. In the single-cell RNA sequencing experiments (Connect_ID_ dataset), we compared the projection profiles of single barcodes for the 52 neurons which had multiple barcodes. We quantified matching by measuring the distance between projection profiles (*i.e*. the cumulative areas between curves). These projection profiles were compared within and across cells (as has been performed across pups in point 5, above). The distance of profiles within single cells was smaller than across cells, demonstrating replicable barcode distributions and further documenting reliability of MAPseq after 14 h of infection.

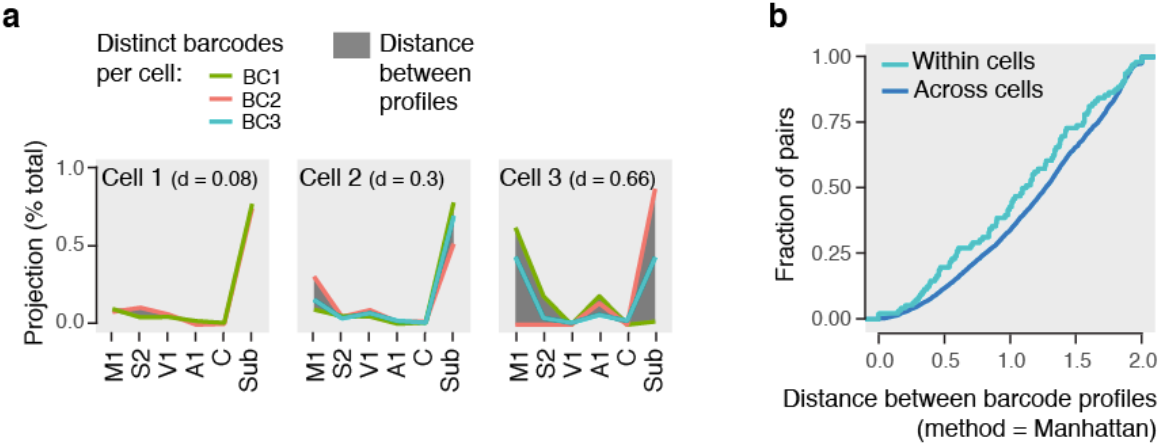
Cumulative distribution of distances between the barcode pairs within and across cells. **a**, Representative projection profiles of cells with multiple barcodes. d: distance between projection profiles. **b**, The shift in the within-cell distribution reflects the higher fraction of closely matched projection profiles, consistent with double infection. Infection time: 14h.

To eliminate potential technical artifacts neurons, we manually examined each of the 52 cells with multiple profiles and eliminated 12 cells displaying high distance between profiles (d > 0.66) with the assumption that they may represent single-cell RNA sequencing doublets.

### Supplementary Note 2. Single-cell RNA sequencing of Sindbis virus-infected cells

Sindbis is an RNA virus, which is likely to affect endogenous transcriptional processes in our experiments. To identify infection-related transcriptional processes and to reduce Sindbis-related transcriptional noise, we applied the following quality control and normalization procedure:

1. We first determined viral load in each single cell, *i.e*. the number of sequenced reads which mapped on the sequence of Sindbis virus^20^.
2. We generated an ordinal regression model to identify genes with the strongest weight in ranking neurons based on their viral load (*i.e*. number of Sindbis reads). This identified genes whose expression was affected by the virus.
3. Ontologies of high-weight genes (FDR < 0.1, *n* = 223 genes) included ribosomal and translation-related processes, consistent with the early stages of hijacking of the cell’s translational machinery by the virus (see table below and **Table S1**). Significantly enriched ontologies assumed to represent reaction to viral infection are listed below; corresponding affected genes present in the dataset were excluded (abs(z) score > 0.5 in the regression model).

4. Gene expression was corrected based on viral load and on the number of expressed genes for each cell using a linear correction^40^, thus mitigating the effect of viral infection. The analyses were performed on the remaining 8419 genes.

| Ontology | FDR |
| --- | --- |
| Inter species interaction between organisms | 2.73 e-22 |
| Mitochondrion | 5.32 e-16 |
| Ribosome | 6.61 e-15 |
| Viral life cycle | 4.25 e-13 |
| Regulation of response to stress | 1.01 e-12 |
| Reactome influenza viral RNA transcription and replication | 3.93 e-12 |
| Response to virus | 8.98 e-12 |
| Defense response | 8.00 e-11 |
| Immune system process | 3.57 e-10 |

**Supplementary Figure 1:**
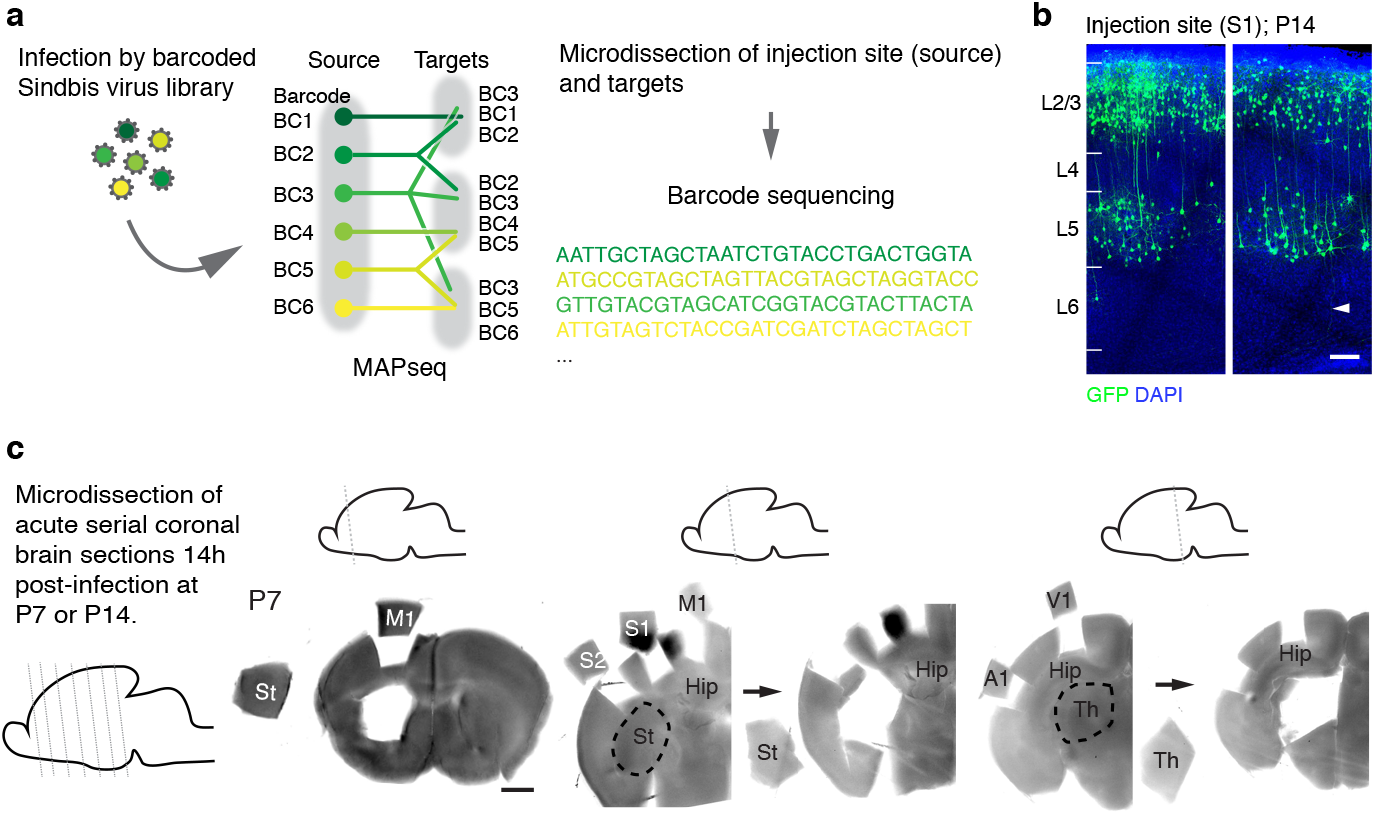
MAPseq experimental procedure. **a**, MAPseq principle. **b**, Infected neurons in S1 expressing Sindbis-GFP 14h after infection. Arrowhead shows an axon labeled with GFP. **c**, Microdissections of injection and target sites. S1, primary somatosensory cortex; M1, primary motor cortex; St, striatum; S2, secondary somatosensory cortex; Hip, hippocampus; V1, primary visual cortex; A1, primary auditory cortex; Th, thalamus. Scale bars, 100 μm (b), 500 μm (c).

**Supplementary Figure 2:**
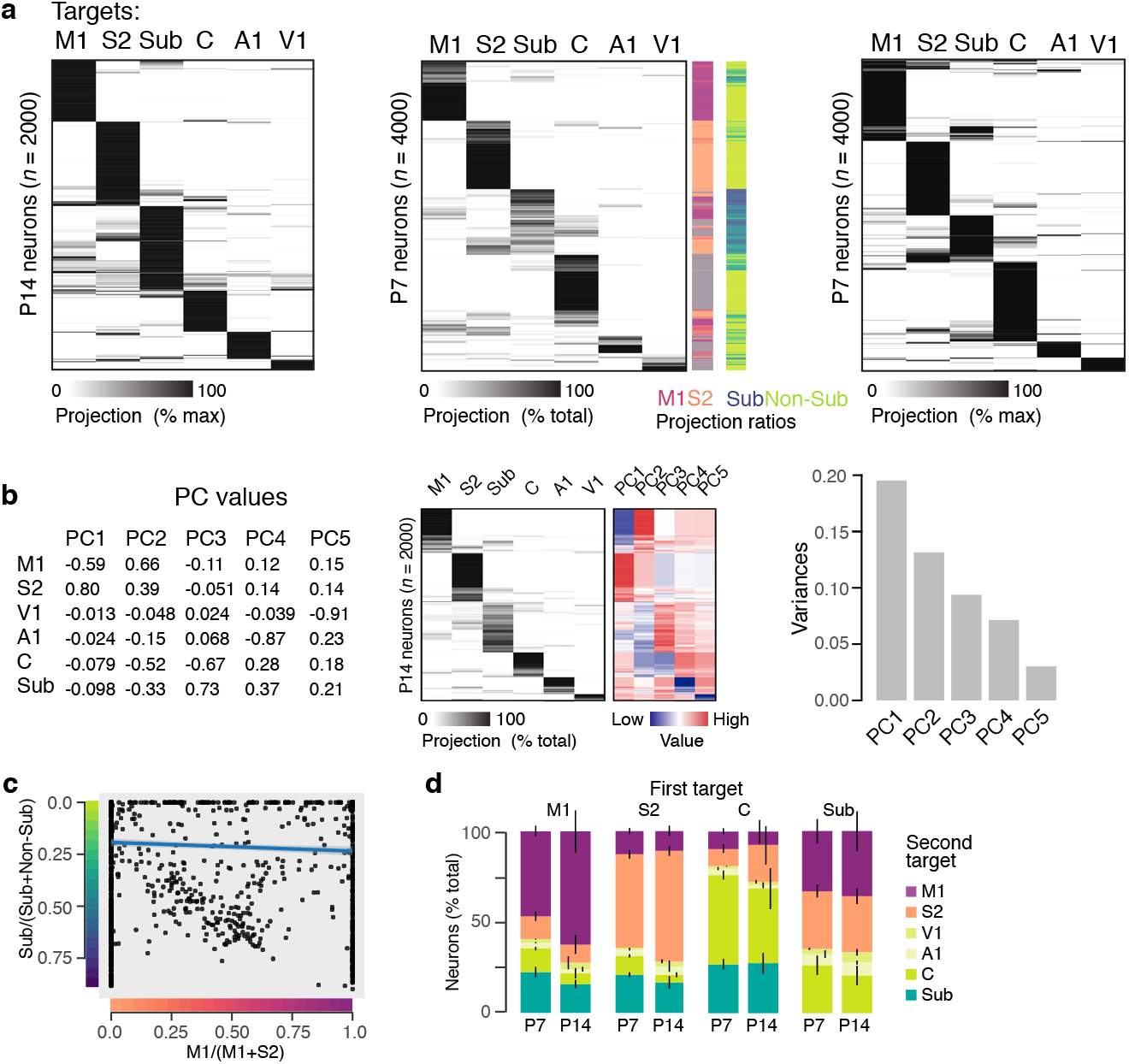
S1 ICPN projections are stable between P7 and P14. **a**, S1 ICPN multiplex projections at P14 (showing data as % max projection) and P7 (showing data as % total projections and % max projection). **b**, Principal component analysis of ICPN projections. Left, mean value of PCs for each target. Center, PC values of single ICPN. Projection heatmap reproduced from Fig. 1b. Right, contribution of each PC showed by their respective variance. **c**, Sub/Non-Sub and M1/S2 ratios of ICPN at P14. **d**, First and second targets of ICPN at P7 and P14. Errors bars, SE; P7: *n* = 4 pups, P14: *n* = 5 pups.

**Supplementary Fig. 3:**
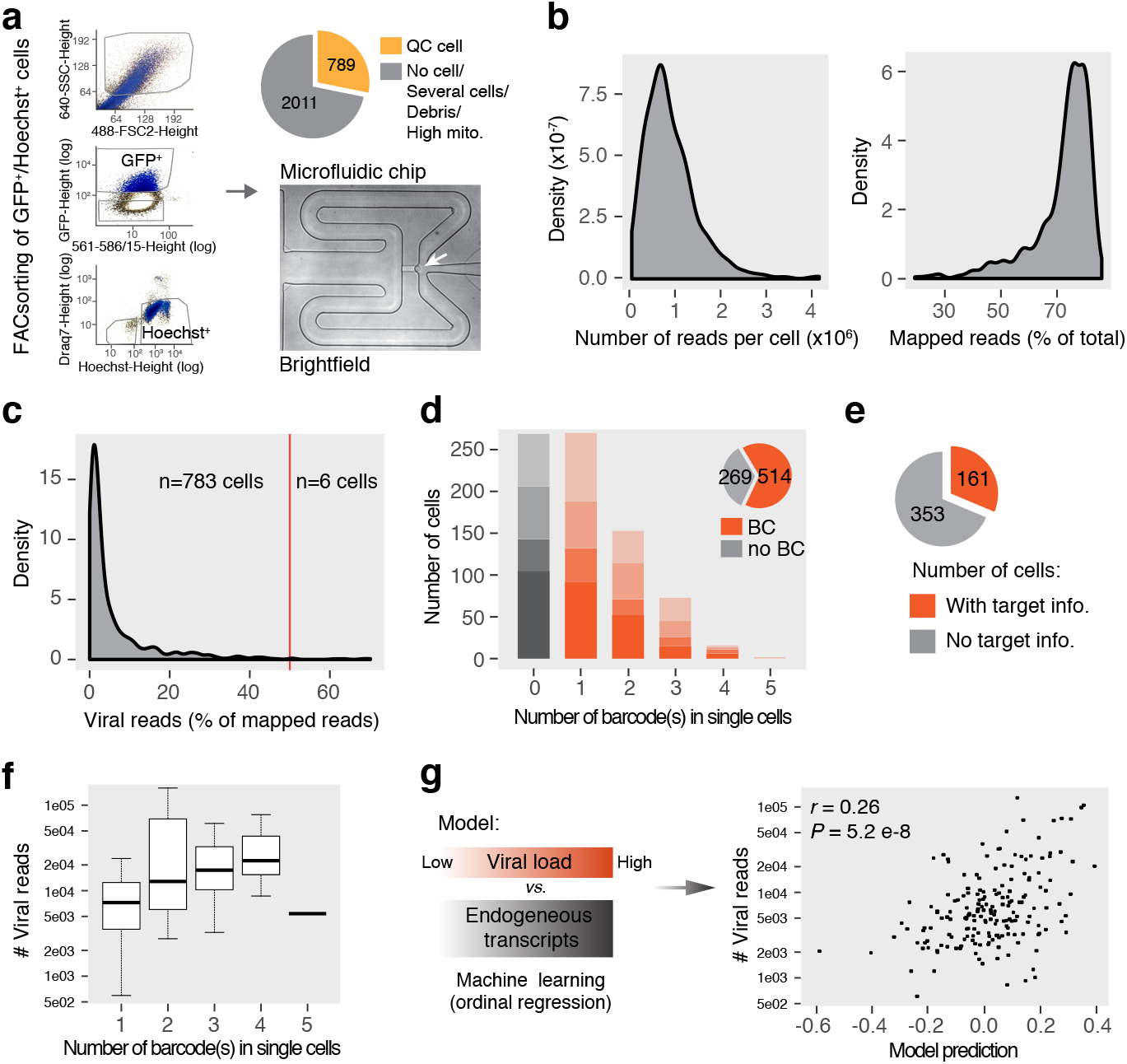
Quality controls of single neurons collected 14 hours after Sindbis infection. **a**, Fluorescence activated cell (FAC) sorting of Sindbis-GFP^+^/Hoechst^+^ neurons (left), capture and quality controls of single cells in microfluidic wells using brightfield imaging (right). Wells with no cell, several cells, debris or with high mitochondrial reads (mito) were excluded, yielding to 789 quality-controlled (QC) cells. **b**, Number of reads per cell (left) and percentage of mapped reads per cell (right). **c**, Percentage of viral reads. Cells with more than 50% of viral reads were excluded from analyses (*n* = 6 cells). **d**, Number of barcode(s) (BC) retrieved in single cell somata after single-cell RNA sequencing (see Methods). Shades of colors: 4 distinct experiments. **e**, Proportion of cells with retrieved BC in the target(s). *n* = 161 cells had information about target(s). **f**, Viral reads increased with the number of barcode(s) per cell. Error bars, SD. **g**, Ordinal regression model identifies genes with the strongest weight in ranking neurons based on their viral load (*i.e*. number of viral reads).

**Supplementary Fig. 4:**
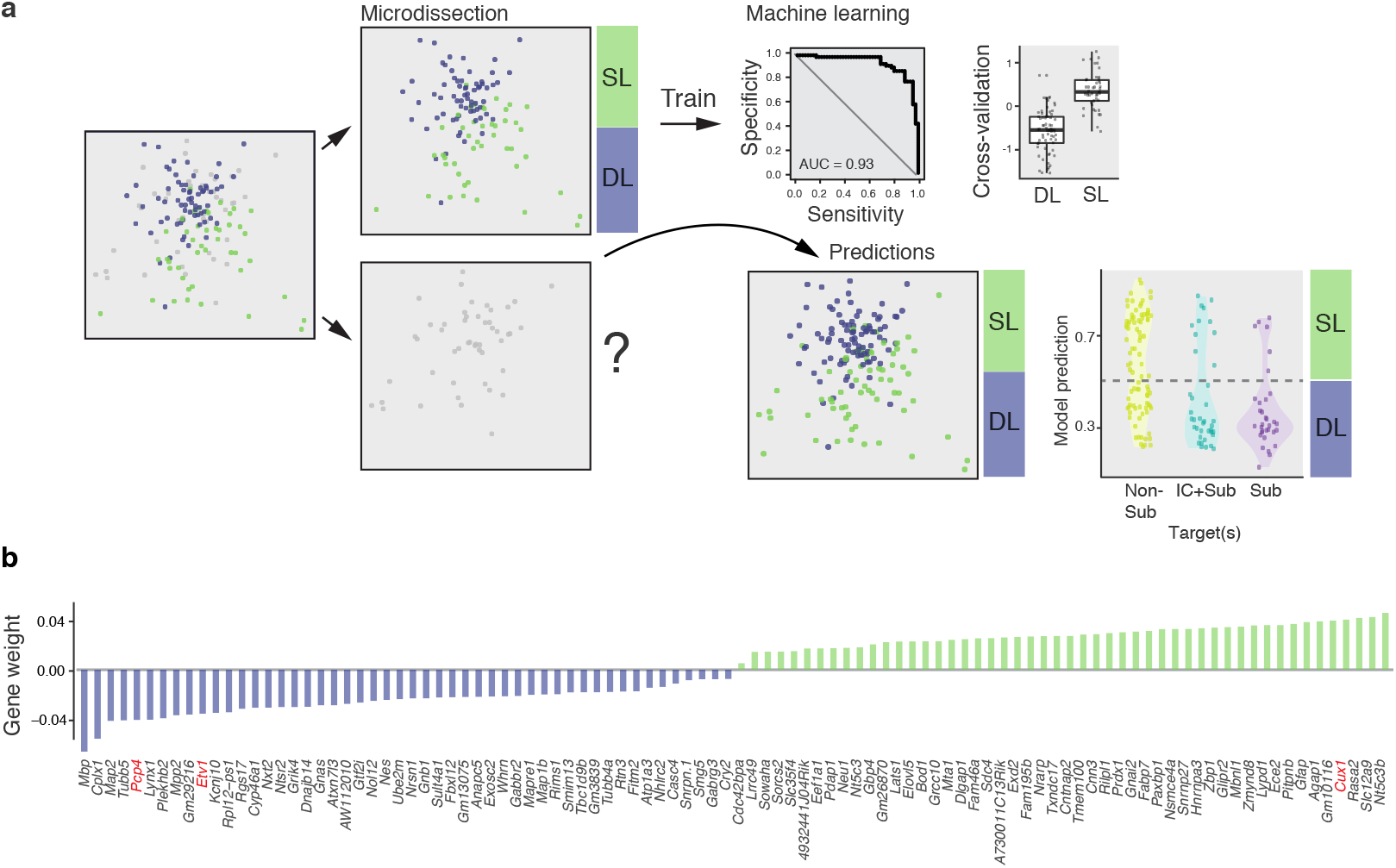
Definition of neuron laminar position using machine learning. **a**, Seed cells from microdissected S1 layers were used to train a linear regression model predicting laminar position. This approach defined best genes distinguishing superficial (SL) from deep (DL) layer neurons with a high confidence showed by the area under roc curve (AUC = 0.93) and the cross-validation (see Methods). Error bars, SD. The model was then used to predict the laminar position of neurons with no layer information. Bottom right, predicted layer position of neurons projecting intracortically (IC) and/or subcortically (Sub). **b**, Genes identified by the model, with their corresponding weight, to distinguish SL *vs*. DL position. In red, are indicated previously reported layer markers.

**Supplementary Fig. 5:**
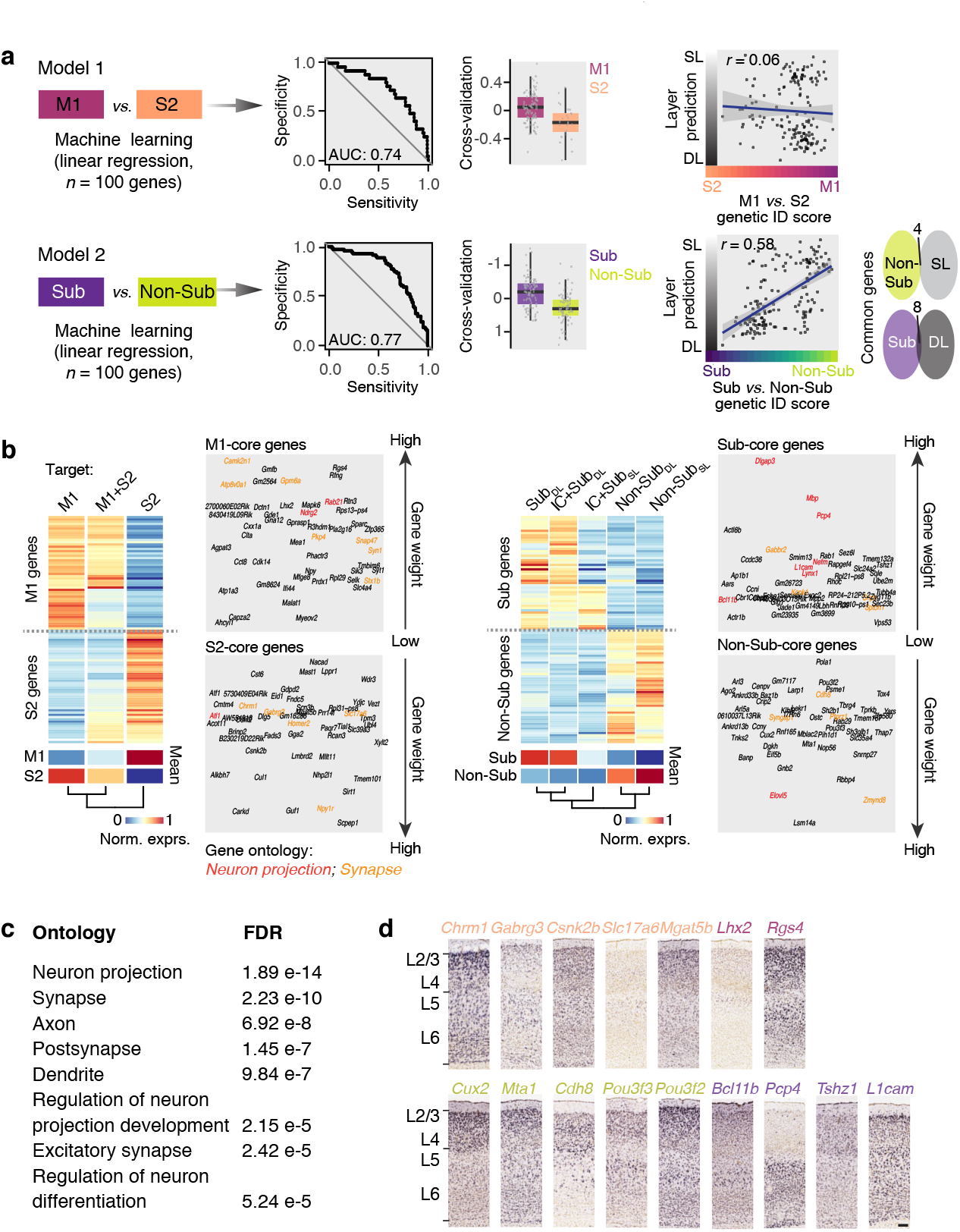
Machine learning identifies genes predicting specific projection motifs. **a**, Two model trainings to predict projection to M1 *vs*. S2, or Sub *vs*. Non-Sub, using linear regression with top 50 genes of each group (left). Model efficiencies are shown with area under roc curve (AUC) and cross-validations (middle). Error bars, SD. Correlation between layer and projection prediction for each cell (right). Number of common genes between the Sub *vs*. Non-Sub projection and layer position models (bottom right), **b**, Expression heatmap of the 100 genes of each model, by distinct projecting neuron classes. Weight of each gene in each model. Left, model 1 ? Right, model 2, IC: intracortical. **c,** Top ontologies of the discriminative genes defined by both models (n = 200 genes), **d,** *In situ* hybridizations of genes referenced in the Allen Brain Atlas (developing brain, P14). Scale bar, 100 μm.

